# Deep conservation of response element variants regulating plant hormonal responses

**DOI:** 10.1101/544684

**Authors:** Lieberman-Lazarovich Michal, Yahav Chen, Israeli Alon, Efroni Idan

## Abstract

Phytohormones regulate many aspects of plant life by activating transcription factors (TF) that bind sequence-specific response elements (RE) in regulatory regions of target genes. Despite their short length, REs are degenerate with a core of just 3-4bps. This degeneracy is paradoxical, as it reduces specificity, and REs are extremely common in the genome. To study whether RE degeneracy might serve a biological function we developed an algorithm for the detection of regulatory sequence conservation, and applied it to phytohormones REs in 45 angiosperms. Surprisingly, we found that specific RE variants are highly conserved in core hormone response genes. Experimental evidence showed that specific variants act to regulate the magnitude and spatial profile of hormonal response in *Arabidopsis* and tomato. Our results suggest that hormone-regulated TFs bind a spectrum of REs, each coding for a distinct transcriptional response profile. Our approach is applicable for precise editing and rational promoter design.

## Introduction

Phytohormones activate coordinated transcriptional responses to regulate plant development and physiology. These responses are mediated by transcription factor (TF) families, such as *AUXIN RESPONSE FACTOR, B-CLASS RESPONSE REGULATORS* and *ABF*, which mediate the response to auxin, cytokinin and abscisic acid, respectively^1–3^. TFs control the expression of their target genes by binding specific nucleotide sequences, termed response elements (REs), in the regulatory region of those genes. The sequence preference of each TF is commonly portrayed as a Position Weight Matrix (PWM), or a DNA motif, which presents the contribution of each nucleotide position to the TF binding. Motifs for the main hormone response regulating TF were derived based on biochemical affinity methods and on enriched sequences in the promoters of genes induced by hormone application^4,5^. These motifs represent a consensus of many sequences and are composed of a short core of invariant nucleotides and several variant nucleotides which identity has little effect on TF binding. The biological role of these variant nucleotides in mediating TF-DNA binding and transcriptional response is unclear. While these motifs were determined by a consensus sequence, hormone-induced transcriptional response of individual genes varies in term of magnitude, tissue-specificity and other conditions. This gene-specific information is lost when considering only the commonalities between many regulatory sequences. Indeed, when attention was paid to the different response profiles of different genes, more specific motifs and flanking sequences were identified^6,7^.

Amongst plant hormones, auxin serves a key role in plant biology and its core response pathway is conserved from basal to higher plants^8^. In its canonical transcriptional pathway, auxin induces the activation of a family of Auxin Response Factor (ARF) transcriptional factors, that bind the auxin response element (auxRE)^1^. The auxRE was first defined as the hexamer TGTCTC, and DR5, a synthetic promoter comprised of eight direct repeats of this motif, could grant auxin-responsiveness to downstream genes^9–12^. Biochemical characterization has shown that ARF proteins bind a more permissive TGTC<u>NN</u> motif^13,14^. Similarly to auxin, the hormone cytokinin activates target genes and increases chromatin accessibility through B-class Response Regulators (RR) that bind the degenerate RE <u>D</u>GAT<u>YN</u> (cytRE; D = A,G,T; Y=C,T)^15–18^. Abscisic acid regulates the transcription of downstream genes via TFs, such as ABF, which bind the degenerate RE <u>B</u>ACGTG<u>K</u> (abRE; B = C,G,T; K=G,T)^3,19,20^.

Recently, it was shown that modification of the DR5 element from TGTC<u>TC</u> to TGTC<u>GG</u> produced a stronger response to auxin^13,21^, suggesting that the variant nucleotides in the motif may affect the transcriptional response profile. However, it is unknown whether these nucleotide variants serve any function in a native sequence context or in the plant itself. It is generally accepted that the existence of purifying selection, evidenced by high sequence conservation, is a strong indicator of functionality^22^. Detecting conservation for specific RE variations is not trivial as rapid structural changes and the fact that RE exists in multiple copies prevents the alignment of regulatory regions. Further, due to common gene duplications in plants, ortholog assignment in divergent species in often uncertain. Indeed, plant studies of regulatory region conservation have restricted themselves to individual loci^11,23^, or focused on ecotypes or a limited number of species^24,25^.

Existing methods for detection of binding site conservation are geared towards the de-novo identification of motifs^26–30^. To enable the detection of variant sequence conservation, we developed Conservation of Motif Variants (CoMoVa), an alignment-free method for detection of conserved small degenerate positions in known motifs over broad evolutionary distances. We applied CoMoVa to detect conservation of hormonal REs in 45 different angiosperms, separated by approximately 150 million years^31^ and identified highly conserved RE variants in core hormonal response genes. We then show that variants of the auxRE can fine tune the transcriptional response to the hormone.

## Results

### CoMoVa - an algorithm for the identification of RE variant conservation

To test the conservation of variant nucleotides in binding motifs, we first focused on the auxRE. Due to the high enrichment of these RE next to the transcription start site^1,32^ (TSS), we analyzed the sequences 500bp upstream and 100bp downstream to the TSS for all protein-coding genes in 45 angiosperms (Fig. 1A; Supplemental Table S1). A mean 91±7% of genes carried at least one degenerate TGTC<u>NN</u> auxRE, and among those, there were 3.54±0.32 auxREs per gene. To determine possible motif conservation, we first identified candidate orthologs. Using the well annotated *Arabidopsis* genome as a reference, a reciprocal blast was performed to identify up to eight candidate orthologs from each species (Supplemental Table S2). The candidates with the most similar set of motif variants were selected as the likely ortholog. Following ortholog assignment, the auxRE variants for each gene were arranged on a predefined species tree representing known phylogenetic relationships (Fig. 1A-D). This analysis could identify genes with little to no conservation of the variant sequences (Fig. 1B), some conservation (Fig. 1C), or high levels of conservation (Fig. 1D). To quantify the conservation level, maximum parsimony optimization was applied to compute the number of motif changes (gain or loss) for each variant along the tree. The gene and motif variant conservation score was defined as the number of orthologs/species carrying the promoter with a particular variant, minus the number of changes in the tree (multiple repeats of the same variant were counted as one). This metric ranged between 0 to 45, representing no to complete conservation, respectively (Fig. 1B-D). As the distribution of dinucleotides is genome-specific^33^ (Supplemental Fig. 1A), background rates were determined empirically, by calculating the conservation score distribution for the two variant nucleotides 8bp upstream to the core sequence (<u>NN</u>nnnnnnnnTGTC; Supplemental Fig. 1B). The resulting distribution of conservation scores was used to compute p-values for significant conservation of variant nucleotide in the auxRE motif.

**Figure 1.**
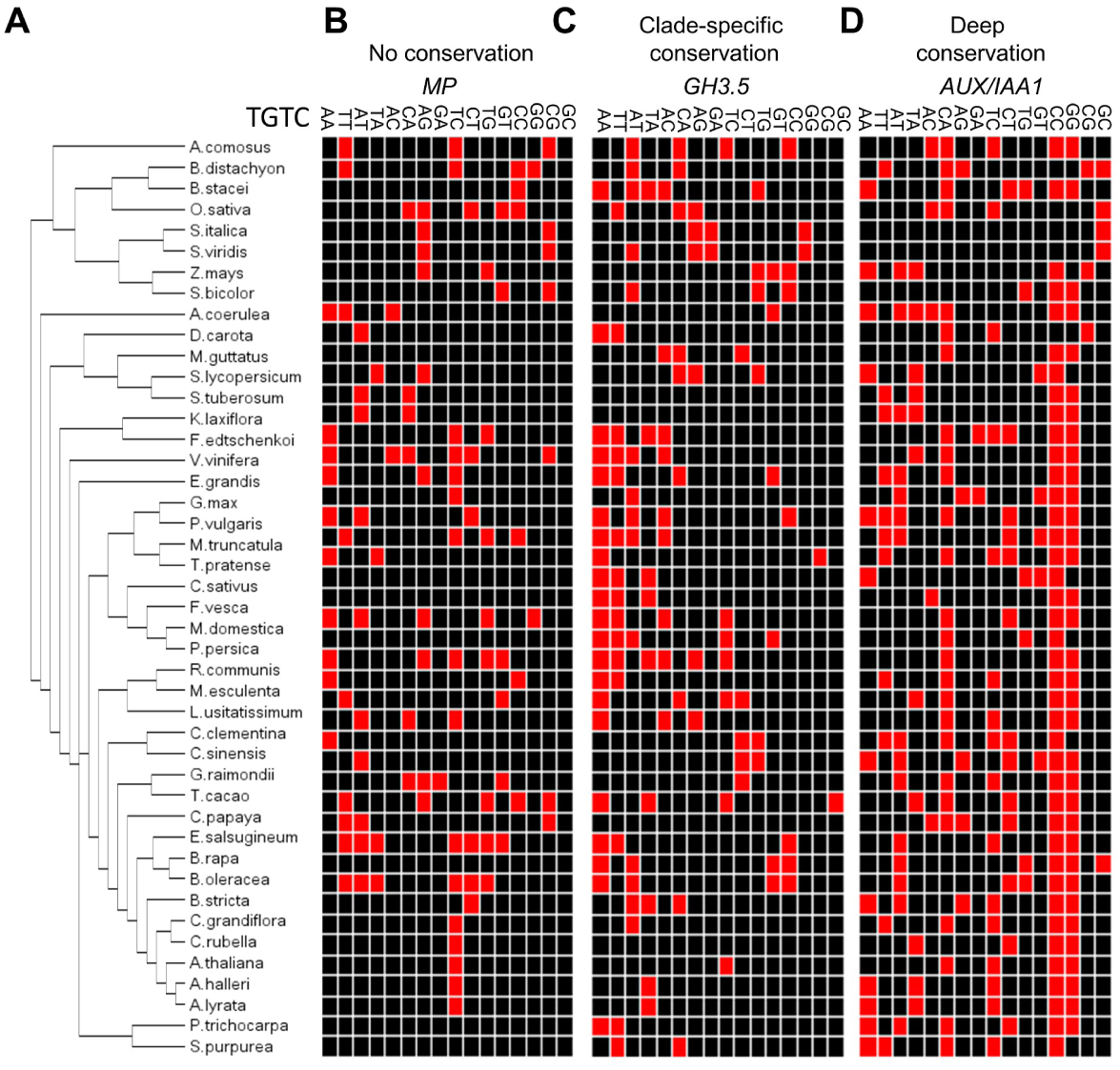
Algorithm for identification of DNA motif variant conservation. **A)** Phylogenetic tree of the species used in this study. B-D) The occurrences of particular motif variants in orthologs of MP (B) GH3.5 (C) and AUX/IAA1 (D). Red indicates the existence of the motif in the gene promoter.

### auxRE motif variant conservation defines the core auxin response

AuxREs are extremely common in the genome, but surprisingly, CoMoVa detected highly significant conservation for a set of just 112 genes. There was a marked bias in the conserved variants, with over 80% of the genes carrying conserved CC, GG, GA, TC, CA, AC or AT motifs (Fig. 2A; Supplemental Table S3). GO-term enrichment analysis of genes with conserved variants revealed highly significant enrichment for response to auxin, auxin homeostasis and auxin activated signaling pathway genes (Fig. 2B). As a control, GO analysis for all genes with auxRE yielded no enrichment for auxin-related terms (Supplemental Fig. 2A). The number of auxRE motifs per gene for genes with conserved auxRE motifs was 3.28±0.49, which was not significantly different that the number for all genes. Interestingly, there was functional separation between genes enriched for different variants. More specifically, genes associated with a conserved TGTC<u>GG</u> variant were enriched for cell wall synthesis functions, genes with conserved TGTC<u>CC</u> were associated with auxin response, while TGTC<u>GA</u> variants were ascribed cellular differentiation roles. Examination of the most highly conserved genes showed that the TGT<u>CC</u> motif variant was conserved in promoters of four AUX/IAA genes (*IAA1/IAA2/IAA3/IAA4*) and in three GH3.3 genes (Supplemental Fig. 2B). Both gene families function in the core auxin response and were shown to be auxin-induced in multiple species^34^. Amongst the genes containing a conserved TGTC<u>GG</u> variant, were the cell wall modifiers GH9B7 and GH9B5, previously shown to act downstream of auxin in lateral root formation,^35^ as well as two EXPANSIN two mannose biosynthesis genes (Supplemental Figure 2B). TGTC<u>CC</u> was also found in association with the transcriptional factor WUSCHEL, which acts to maintain the growth of the shoot apical meristem ^36^ and VND2, which regulates xylem formation ^37^.

**Figure 2.**
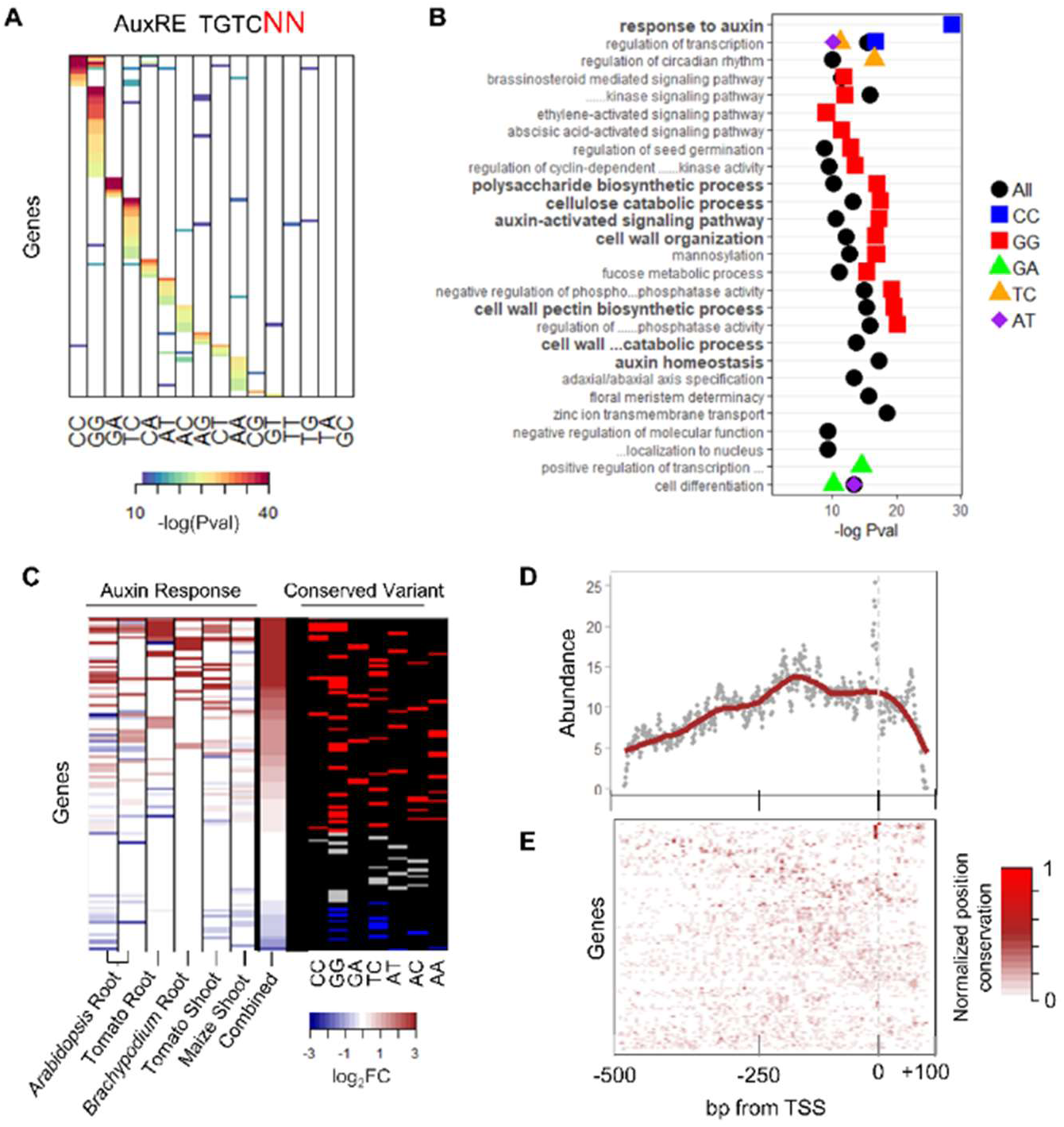
Conservation of auxRE variants. **A)** Genes and motif variants showing significant conservation. Each column represents a specific dinucleotide variant of the TGTC<u>NN</u> motif. **B)** GO term enrichment of genes with a conserved auxRE motif variant. **C)** Auxin responsiveness of genes with conserved auxRE variants in different tissues and species. **D)** Mean abundance of conserved motifs in each position relative to the TSS for all 45 angiosperms. **E)** Conservation of the auxRE motif position for each gene for the 45 angiosperms species. High values (red) represent high conservation of the motif position.

Three of the highly conserved variants (TGTC<u>GG</u>/<u>CC</u>/<u>TC</u>) were previously identified as enriched in sets of auxin-responsive genes in *Arabidopsis*^5,7,9,32^. To test for auxin responsiveness of genes across multiple species, we complied five rapid auxin response-related transcriptional activity datasets in *Arabidopsis*, maize and *Brachypodium*. To add additional dicots, we also generated RNA-Seq profiles of tomato (*Solanum lycoperiscum*) roots and shoots at 3h following auxin application. While auxin response differed between organs and species, most of the genes with conserved variants exhibited auxin response in at least one sample (Fig. 2C).

We next sought to determine whether the position of the auxRE relative to the TSS is also conserved. Despite large differences in the length of intergenic regions between the species, auxREs were enriched in the +50 to −250bp region relative to the TSS across all tested angiosperms (Fig. 2D). Furthermore, for only a subset of genes, we could identify high conservation of the position of the auxRE relative to the TSS (Fig. 2E). Overall, the conservation of specific variants and the consistent biological role of genes with conserved auxRE motifs suggests that CoMoVa identified an ancient regulatory module acting downstream of auxin, guiding negative feedback and cell wall modifications. The conservation of specific auxRE variants across vast evolutionary distances suggests strong selection pressure on the variable dinucleotides, indicating their identity is important for gene function.

### Motif variant conservation defines core response for cytokinin and abscisic acid

If the conservation of variable nucleotides in promoters of core response genes is a general phenomenon, we hypothesized that applying the same algorithm to other hormonal RE will identify core target genes for these hormones. To test this hypothesis, we applied CoMoVa to characterize the conservation of the cytRE (<u>D</u>GAT<u>YN</u>), which mediates the cytokinin response and appears in 96.4±0.6% of the genes in all species. In agreement with our hypothesis, CoMoVa identified highly conserved <u>A</u>GAT<u>TT</u> and <u>G</u>GAT<u>TT</u> variants in seven of the ten A-class RRs (Fig. 3A-B). These negative feedback regulators of the cytokinin response play a role in the core cytokinin response in multiple species^2^.

**Figure 3.**
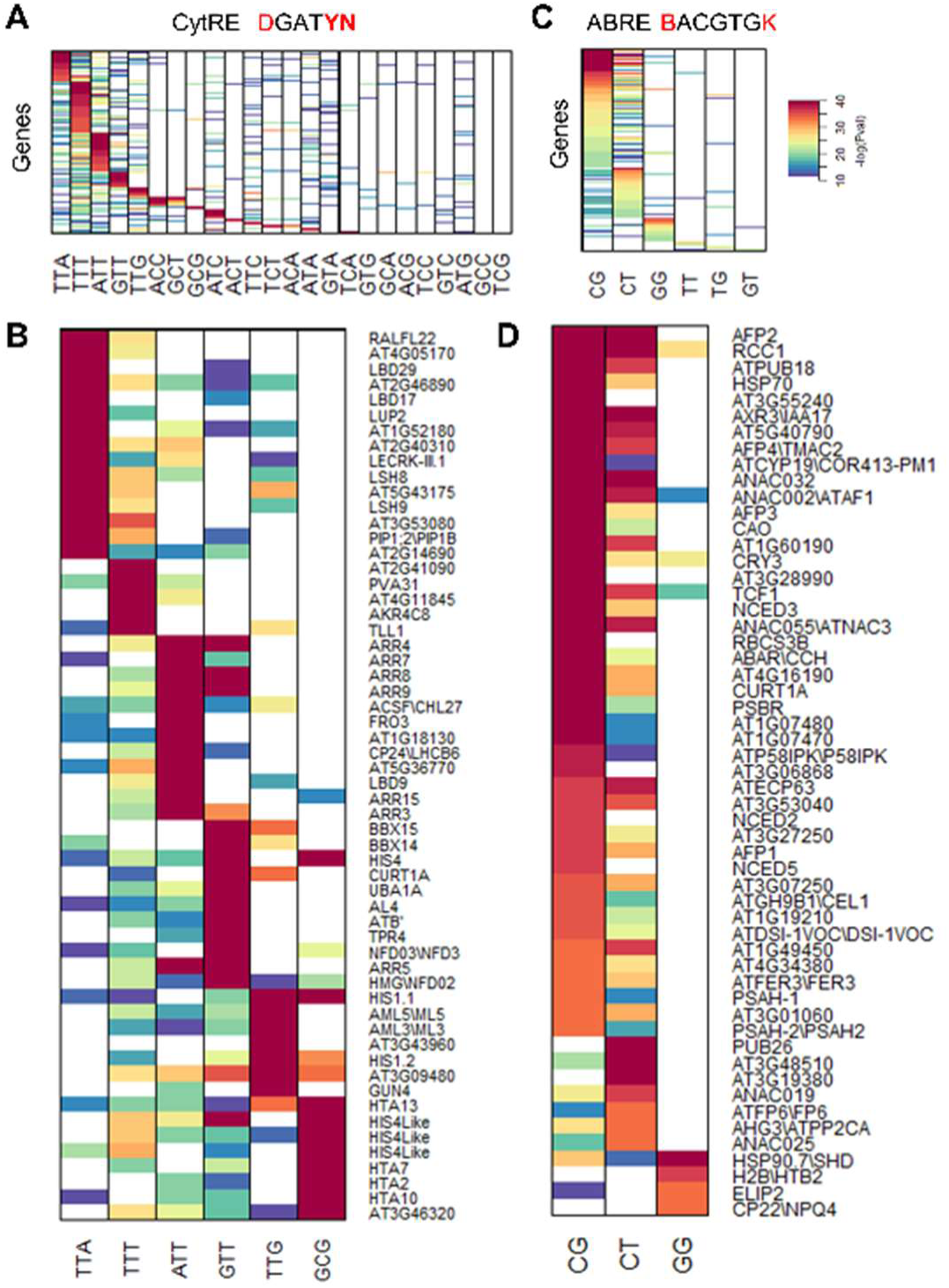
Conservation of cytRE and abRE variants. **A-D)** Conservation of cytRE (A-B) and abRE (C-D) motif variants, all showing significant conservation (A,C) or just the genes with most highly conserved motifs (B,D).

Overall, conservation of 10 motifs (out of the possible 24) was detected in 537 genes which were enriched for cell division regulators, histones and histone modifiers (Supplemental Fig. 3A; Supplemental Table S4). Multiple *HISTONE H4* and *HISTONE H2A* genes had a strongly conserved <u>G</u>GAT<u>CG</u> motif, while the G1-S regulators CYCA3;4 and CYCD4;2 had conserved <u>A</u>GAT<u>TT</u> and <u>T</u>GAT<u>TT</u> motifs (Fig. 3B). Unlike the auxRE motif, no conservation of cytRE motif position relative to the TSS was detected (Supplemental Fig. 3B).

As an additional control, we tested the conservation of the abRE motif, which mediates the ABA response and appeared in 21±5% of all genes in all species. CoMoVa identified motif conservation (<u>C</u>ACGTG<u>G</u>) in known ABA-inducible genes, such as the ABA biosynthetic enzymes NCED2/3/5/9^38,39^ and members of the ABI Five Binding Protein (AFP) family^40^ (Fig. 3C-D). GO term analysis of the 238 genes with conserved abRE motifs showed enrichment for ABA response, regulation of light responses, response to cold and seed germination (Supplemental Fig. 4A; Supplemental Table S5). Moreover, despite large differences in intergenic region length between the species, conserved abRE variants were highly enriched at −150bp from the TSS, suggesting that this position is important for the RE function (Supplemental Fig. 4B).

Overall, motif variant conservation appears to be prevalent and core hormonal response targets are enriched with specific variants, suggesting that this evolutionary signature can be used to identify core sets of TF targets.

### Deep conservation of motif flanking sequences

Active TF binding sites are characterized by sequence conservation in regions flanking the core binding site^41^. Indeed, promoter mutagenesis has shown that nucleotides flanking the auxRE are critical for generating proper auxin-response^11,12,42^. We therefore sought to identify conservation in regions flanking the auxREs. To this end, the flanking 20bp up- and downstream for each ortholog group were aligned, centering on the conserved motif. A PWM was calculated and the information content for each position served as a measure of conservation at a specific position. As predicted, additional conserved nucleotides were identified in the vicinity of the auxRE (Figure 4A; Supplemental Table S6), which, in some cases, were quite long, such as the stem-cell regulator *WUS*, which had an 18-bp conserved region flanking the TGTC<u>CC</u> motif and a TCCCTTTCTA sequence 20bp upstream.

**Figure 4.**
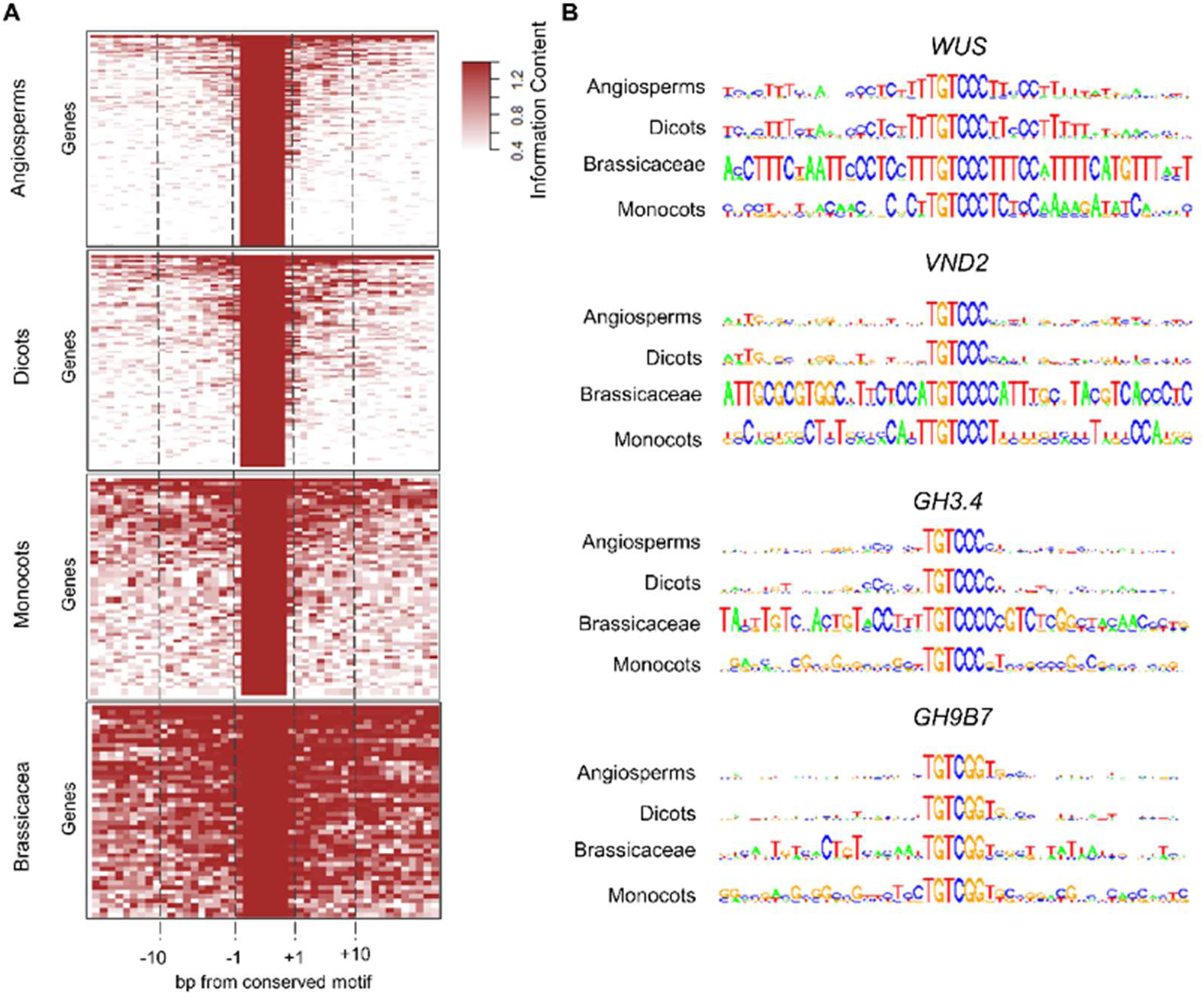
Deep conservation of sequence context for conserved auxRE variants. **A)** Information score values computed from position weight matrix for sequences flanking the conserved auxRE. **B)** Position weight matrix for sequences around the highly conserved auxRE variant.

For many genes, conservation outside the auxRE amongst all angiosperms was weak, suggesting these sequences evolved faster than the variant nucleotides in the motif. As many aspects of gene expression profiles maybe specific to more closely related species, we applied the same algorithm to the dicots, monocots and brassicacea, separately to test whether we can find broader signs of conservation within a shorter evolutionary distances. Significantly higher levels of conservation were detected when the subtrees were analyzed, with clear divergence between monocots and dicots (Fig. 4A-B; Supplemental Table S6). The Brassicaceae family was represented by 10 species in our sample, with around 16 million years between some^43^. Despite this, sequence conservation around the auxRE motifs was very high. The conservation followed gene-specific patterns, but notably, declined with increasing distance from the motif, indicating nucleotides closer to the motif are more highly conserved (Fig. 4A-B). Similarly to the auxRE, sequence conservation for regulatory elements extended beyond the conserved RE for genes containing either cytRE or abRE, and declined away from it (Supplemental Fig. 5A-B; Supplemental Table S7-S8). Overall, our analysis revealed that while the general response to auxin may be defined by the short auxRE motif, the broad sequence context also tends to be conserved. Conservation beyond the motif was more pronounced within plant families, suggesting these sequences may serve family-specific common functionality.

### auxRE motif variants control the transcriptional response profile in synthetic promoters

To identify a possible role of the RE variable nucleotides *in planta*, we generated four versions of the canonical DR5 synthetic promoter, containing different variants of the auxRE motif: the conserved TGTC<u>CC</u>, TGTC<u>TC</u>, and TGTC<u>AC</u>, and the non-conserved TGTC<u>GC</u> (Supplemental Fig. 6A). Promoters were cloned upstream of the fluorescent protein *VENUS, Arabidopsis* root protoplasts were transfected with the construct, treated with auxin for 6h and fluorescence was measured using a flow cytometer. While all four variants responded to auxin in a dose-dependent manner, there was a significant difference in response magnitude, with TGTC<u>CC</u> eliciting the strongest and TGTC<u>AC</u> the weakest response (Fig. 5A). The attenuated response of the TGTC<u>AC</u> motif was unexpected, as *ARFs* bind this sequence *in vitro*^13^. To test whether ARF proteins can activate this motif *in planta*, we overexpressed two class A ARFs, *ARF7* and *ARF8*, together with our reporter. *ARF* overexpression successfully restored the auxin responsiveness of the TGTC<u>AC</u> motif to a similar level as that recorded for TGTC<u>TC</u>, suggesting that *in planta*, a higher ARF concentration can compensate for lower activity associated with specific variant nucleotides (Fig. 5B).

**Figure 5.**
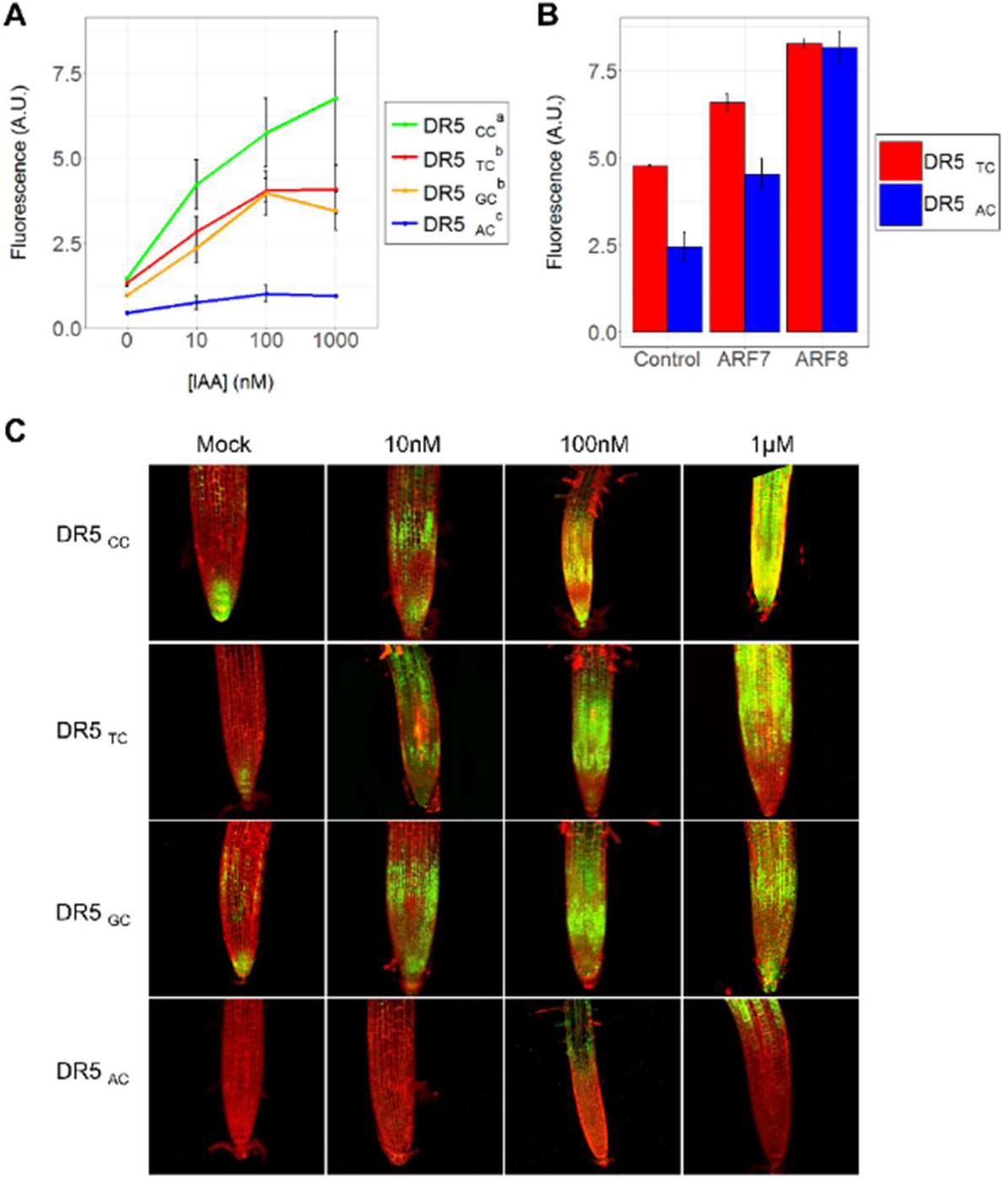
Auxin responsiveness of different auxRE variants. **A-B)** Mean normalized fluorescence of protoplasts transfected with variants of DR5 and treated with auxin for 6h (A) or with DR5 variants and an activating ARF (B). Experiments were performed in triplicates. Error bars represent standard error. (C) Maximal projection of confocal images of roots carrying GFP driven by different auxRE variants. Roots were treated with auxin for 6h.

To test how the variant nucleotides affect transcriptional response in a developmental context, we generated stable-transformed *Arabidopsis* lines of the different DR5 variants fused to GFP, and selected a representative line (out of at least 10 independent lines). At 5 days after germination, seedlings were treated with auxin for 6h. Reporter expression and RNA levels, as measure by qPCR, recapitulated the transient expression results (Fig. 5C; Supplemental Fig. 6B). Interestingly, we observed a difference in the spatial expression pattern, where the response in TGTC<u>AC</u>-bearing lines was restricted to the root elongation zone, as opposite to the almost ubiquitous response of other variants. Notably, the magnitude of transcriptional response mediated by the motif variants did not correlate with its conservation level, suggesting that the conserved motifs are not merely the strongest activators, but mediate a specific response profile.

### Conserved auxRE variant fine-tunes the auxin transcriptional response in native promoters

To test the impact of the motif variants in native promoters, we modified the conserved TGTC<u>CC</u> auxRE motif to the weak-response variant TGTC<u>AC</u> in the promoters of *IAA2* and *IAA4*. Both promoters had a second TGTC<u>CC</u> site with a 5-bp spacer, a configuration preferred by ARF dimers^44,45^. As this pair arrangement was perfectly conserved within the Brassicacea, we mutated both motifs (Fig. 6A-F).

**Figure 6.**
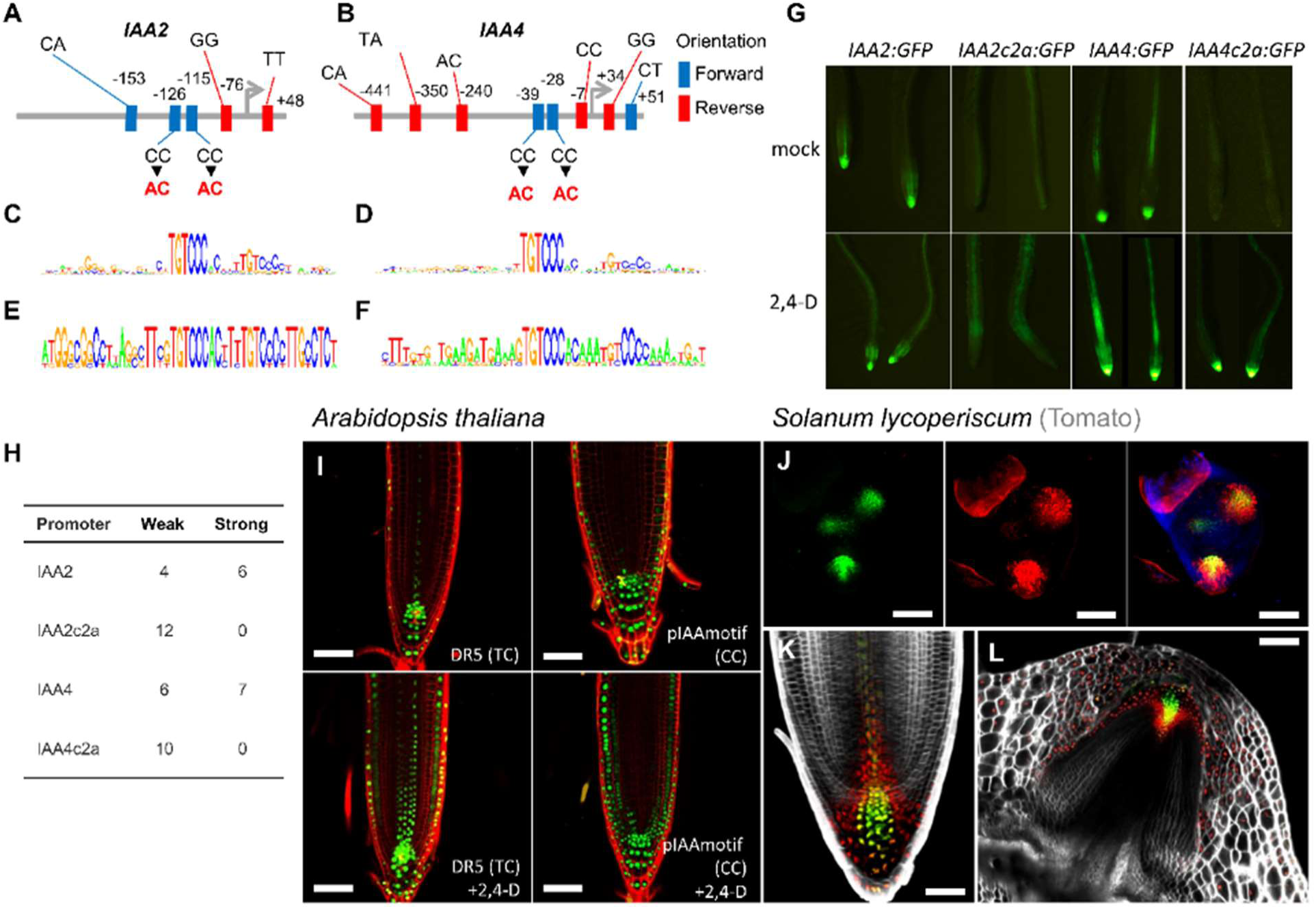
auxRE variants generate predictable auxin response profiles in native and synthetic promoters. **A-B)** Diversity of auxRE motifs at the promoters of *Arabidopsis* IAA2 (A) and IAA4 (B). **C-F)** PWM of the conserved TGTC<u>CC</u> auxRE for IAA2 (C,E) and IAA4 (D,F) in all angiosperms (C-D) or within the Brassicaceae (E-F). **G)** Response of promoters and mutated promoters to auxin. **H)** Number of independent transformation lines exhibiting the same expression as in (G). **I)** Roots of 7 days old *Arabidopsis* plants carrying DR5 and pIAAmotif:mNeonGreen-NLS treated with mock (top) or 1µM 2,4-D (bottom). **J-L)** Shoot meristem (J), main root (K) and stem-borne root (L) of tomato carrying DR5:3xVENUS-N7 (green) and pIAAmotif->opRFP (red). Blue in (J) is chlorophyll autofluorescence. Cell wall stained with Propidium Iodide (I) or SR2200 (K-L).

Roots of *IAA2:GFP* plants exhibited strong fluorescence in the root tip and in the stele, which was upregulated by auxin, consistent with previous reports ^46,47^. In contrast, roots carrying the mutated *pIAA2_cac_:GFP* had weak to no IAA2 expression in the root tip and weak stele expression, with auxin triggering a mild increase in stele expression (Fig. 6G). Roots of *pIAA4_CAC_:GFP* exhibited reduced fluorescence and auxin responsiveness compared to *pIAA4:GFP*, but the response was stronger than *pIAA2_cac_:GFP* (Fig. 6G), possibly due to an additional TGTC<u>CC</u> RE in their promoter (Fig. 6A). To verify that these changes in expression were not mediated by a position effect, we screened multiple insertion lines and found consistent effects of the C to A change in the auxRE (Fig. 6H).

### auxRE conservation enables rational design of synthetic reporters in multiple species

Following our observation that the TGTC<u>CC</u> motif variant in the promoter of the Aux/IAA genes is an important determinant of the magnitude of transcriptional response to auxin, we asked whether we can use this motif to generate a new synthetic reporter for auxin. To maintain conserved position of the motif at −220 to −280bp to the TSS and to avoid introduction of repetitive sequences that are prone to silencing, we cloned three ~50-bp regions of conserved TGTC<u>CC</u>-containing motifs from three different *Arabidopsis* and *Populus trichocarpa* promoters and placed them upstream of a minimal 35S promoter (Supplemental Fig. 7). This promoter (pIAAmotif) was more sensitive to auxin and exhibited a stronger response than DR5 in *Arabidopsis*, but was not as broad as the synthetic TGTC<u>GG</u>-based Dr5v2^21^, adding an intermediate tool for reporting auxin response (Fig. 6I). To determine the evolutionary conservation of these responses, we also generated tomato plants driving mRFP from the same pIAAmotif reporter. Similarly to *Arabidopsis*, the pIAAmotif reporter exhibited overlapping but broader expression domains as compared with the classical DR5 in both roots and shoots (Fig. 6J-L), supporting the notion that the designed high-sensitivity promoters can be used in broad evolutionary contexts.

## Discussion

How promoters encode specific transcriptional responses is still far from being fully understood. The presence of particular DNA motifs contributes to TF binding but cannot explain the complex transcriptional response of the gene. Here, we show that particular variant nucleotides in REs can tune the transcriptional response profile of the promoter and that conservation of the variants nucleotides can be used to identify ancient modules activated by the TF. For example, cytokinins have long been tightly associated with promotion of cell division, but the mechanism by which cytokinins mediate their control of the cell cycle has been elusive^48^. Interestingly, apart from the A-class *RR*, which are known downstream factors of cytokinin, we also identified deep conservation of specific cytREs in promoters of key histones and cell-cycle genes, suggesting that *RRs* may have direct transcriptional control on the cell cycle machinery. While the requirement for a large number of curated genomes limits the approach to identification of highly conserved core functions of the TF, increased availability of plant genomes will enable the detection of conservation between more closely related species and the identification of plant family-specific conserved regulatory regions. Furthermore, the focus of this works was on transcriptional response to hormonal signals, but a degenerate RE was defined to a large set of TF^44,49^ and a similar approach can be applied them.

Motif variant conservation can be used to apply rational promoter design. For example, the recently developed 6xABRE synthetic ABA reporter is based on repeats of the TACGTGTC abRE variant^19^. However, motif conservation analyses showed that this RE is poorly conserved while the ATnnAACACGTGG variant (Fig. 3D; Supplemental Table S8) is strongly conserved in highly responsive NCED genes. It would be interesting to assess whether such variants can be used to generate reporters with different degrees of sensitivity to the hormones. Further, the demonstration that position relative to the TSS is highly conserved for some REs, warrants maintenance of this relative position in synthetic promoters.

Finally, the advent of gene editing techniques has opened possibilities for generation of new alleles in important crops by directed mutagenesis of gene promoters. We propose that RE conservation can be used as a heuristic tool for identification of gene editing targets in order to generate new quantitative alleles for breeding purposes^50^.

## Supporting information

List of genes with conserved abRE motifs

Sequence conservation of sequence flanking conserved auxRE motifs

Sequence conservation of sequence flanking conserved cytRE motifs

Sequence conservation of sequence flanking conserved abRE motifs

List of genomes used in this study

List of genes with conserved auxRE motifs

List of genes with conserved cytRE motifs

Ortholog candidates for all genes in the species used in this study

---

**Supplemental Figure 1.**
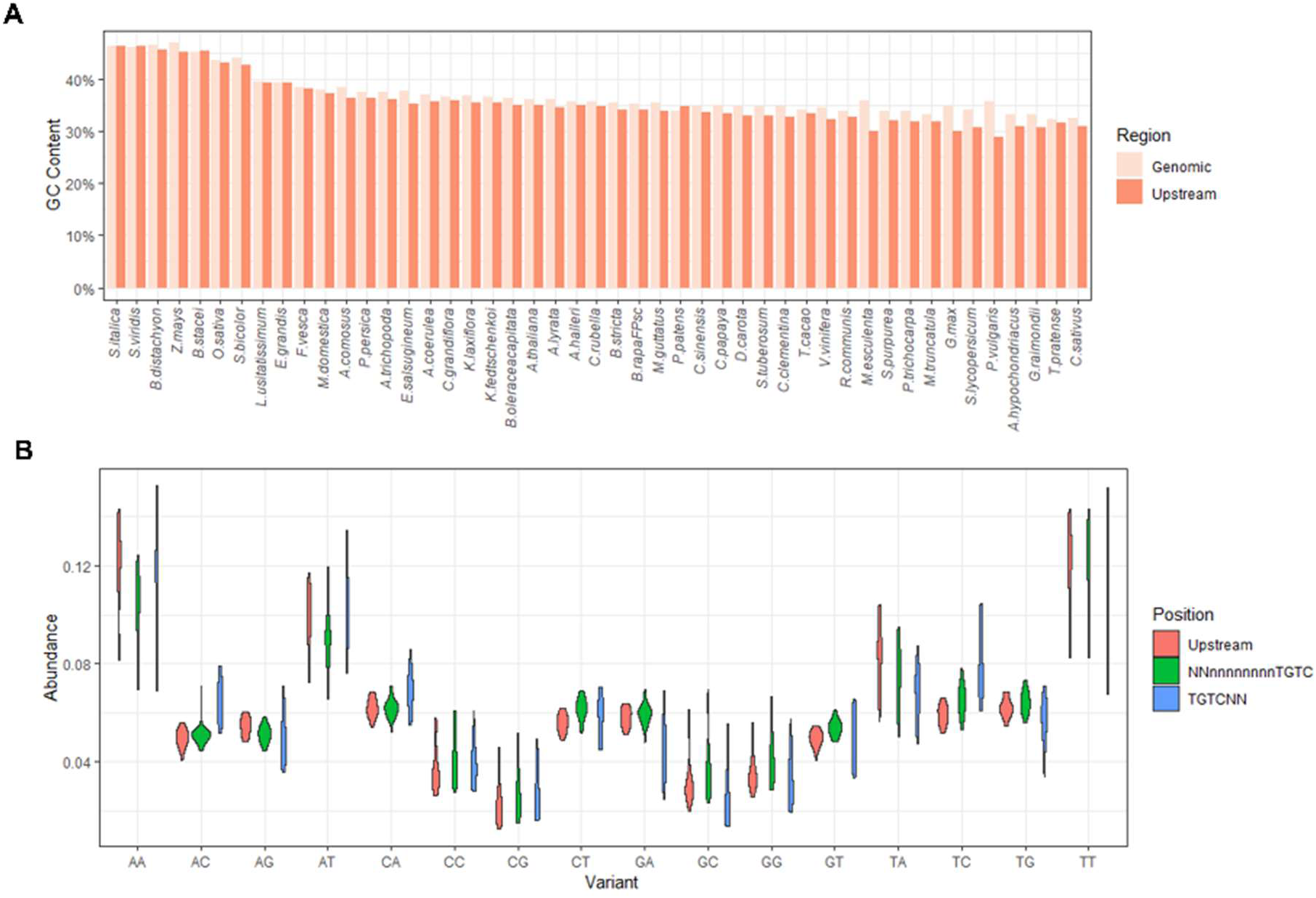
Genome properties of the angiosperms used in this study. **A)** GC content in entire genome and in upstream regions for all genes in the species used in this study. **B)** Distribution of dinucleotides in different genomic regions for all species used in this study.

**Supplemental Figure 2.**
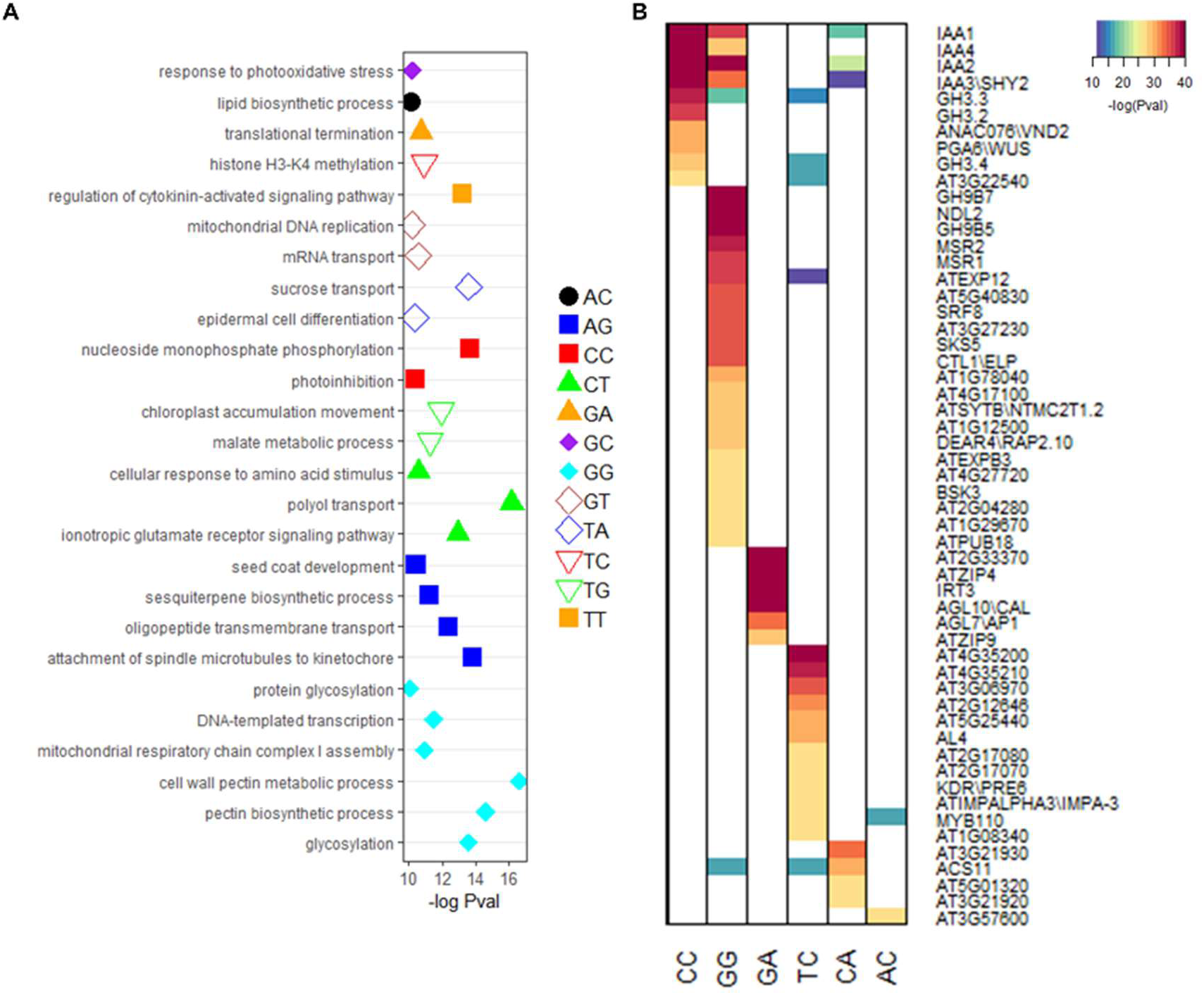
Conservation of auxRE variants. A) GO term enrichment for genes which have the auxRE variant in the promoter region in the *Arabidopsis* genome. B) Genes with most highly conserved auxRE motif.

**Supplemental Figure 3.**
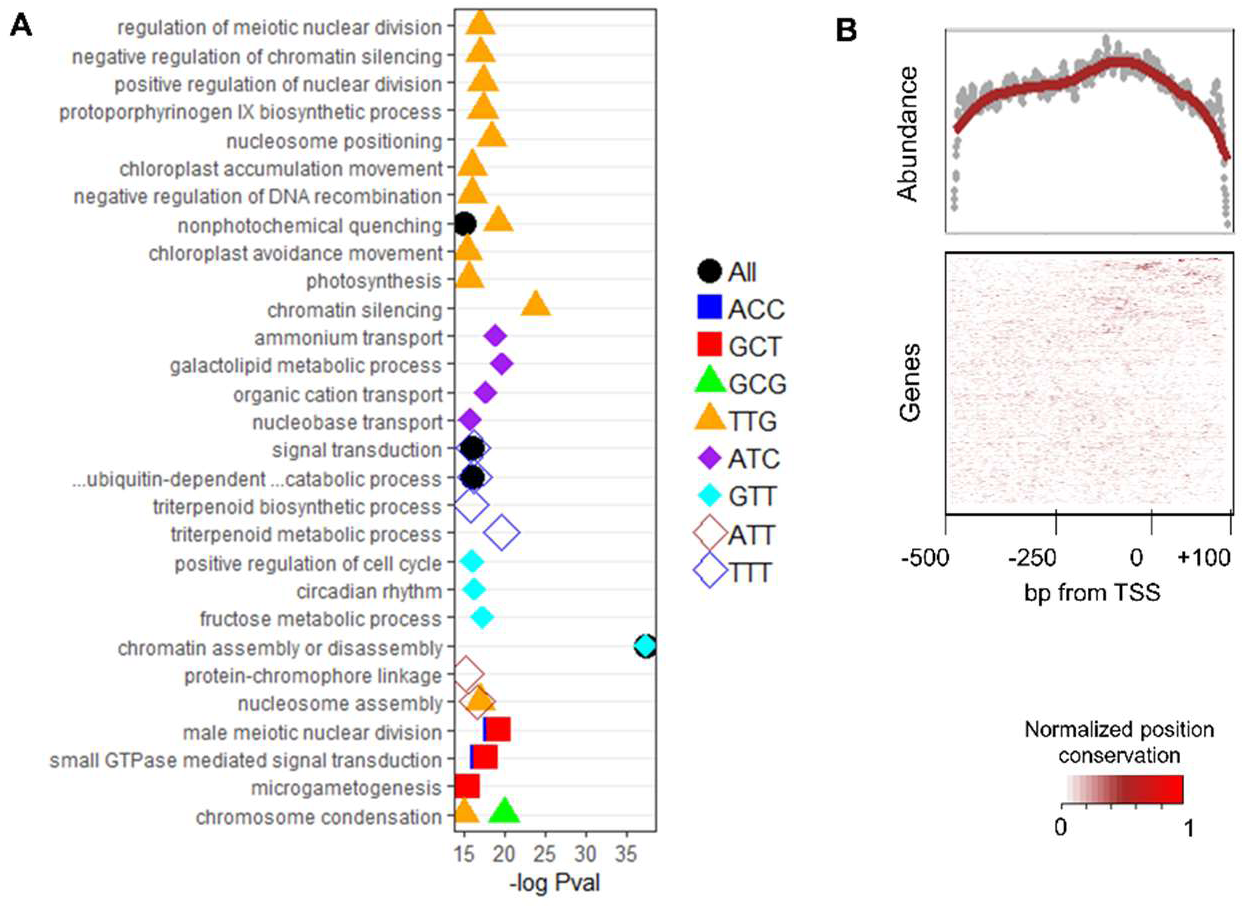
Conservation of cytRE variants. **A)** GO term enrichment of genes with a conserved cytRE motif variant. **B)** Mean abundance of the conserved motif in each position relative to the TSS. **C)** Conservation of the cytRE motif position for each gene. High values (red) represent high conservation of the motif position.

**Supplemental Figure 4.**
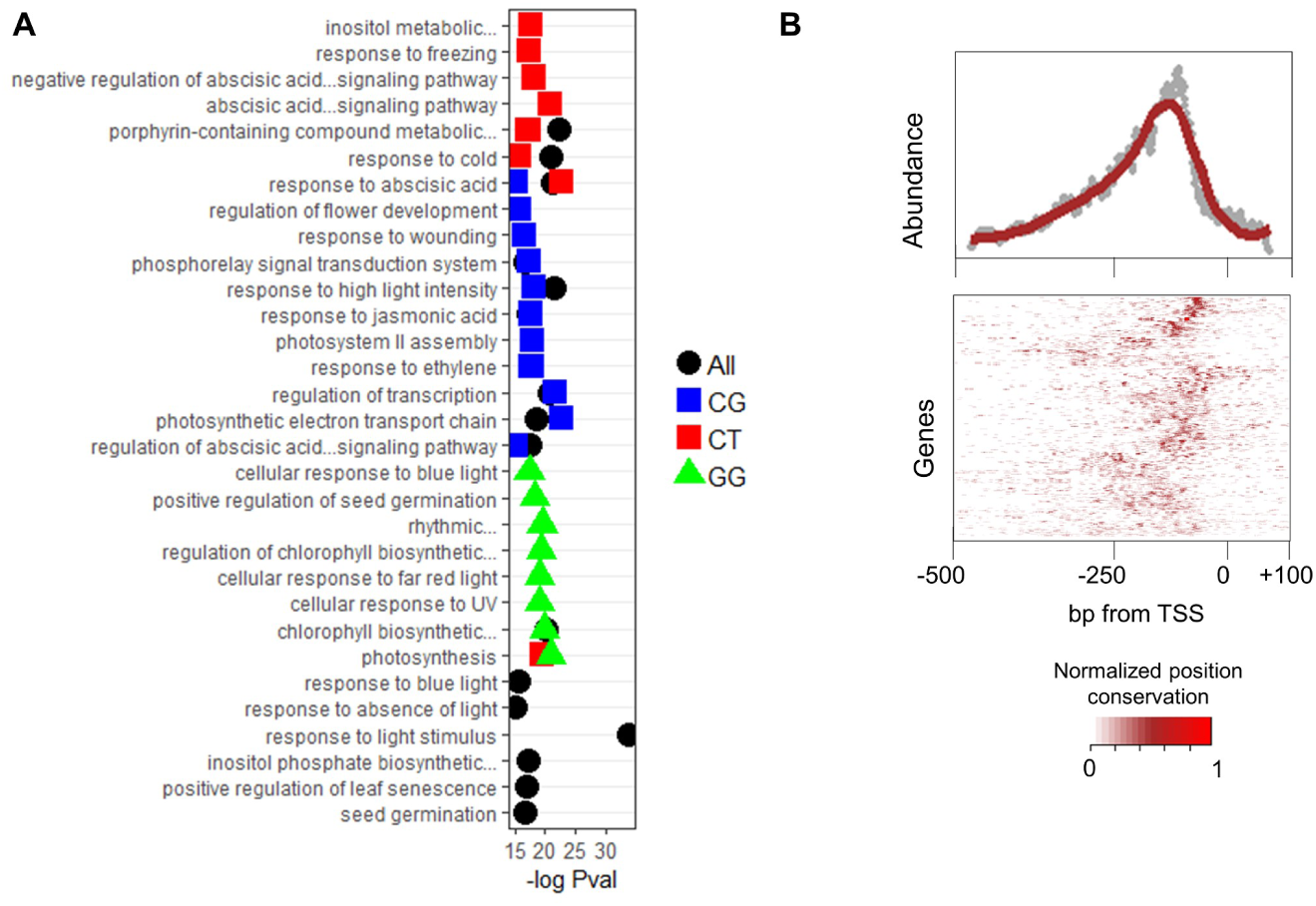
Conservation of abRE variants. **A)** GO term enrichment of genes with a conserved abRE motif variant. **B)** Mean abundance of the conserved motif in each position relative to the TSS. **C)** Conservation of the abRE motif position for each gene. High values (red) represent high conservation of the motif position.

**Supplemental Figure 5.**
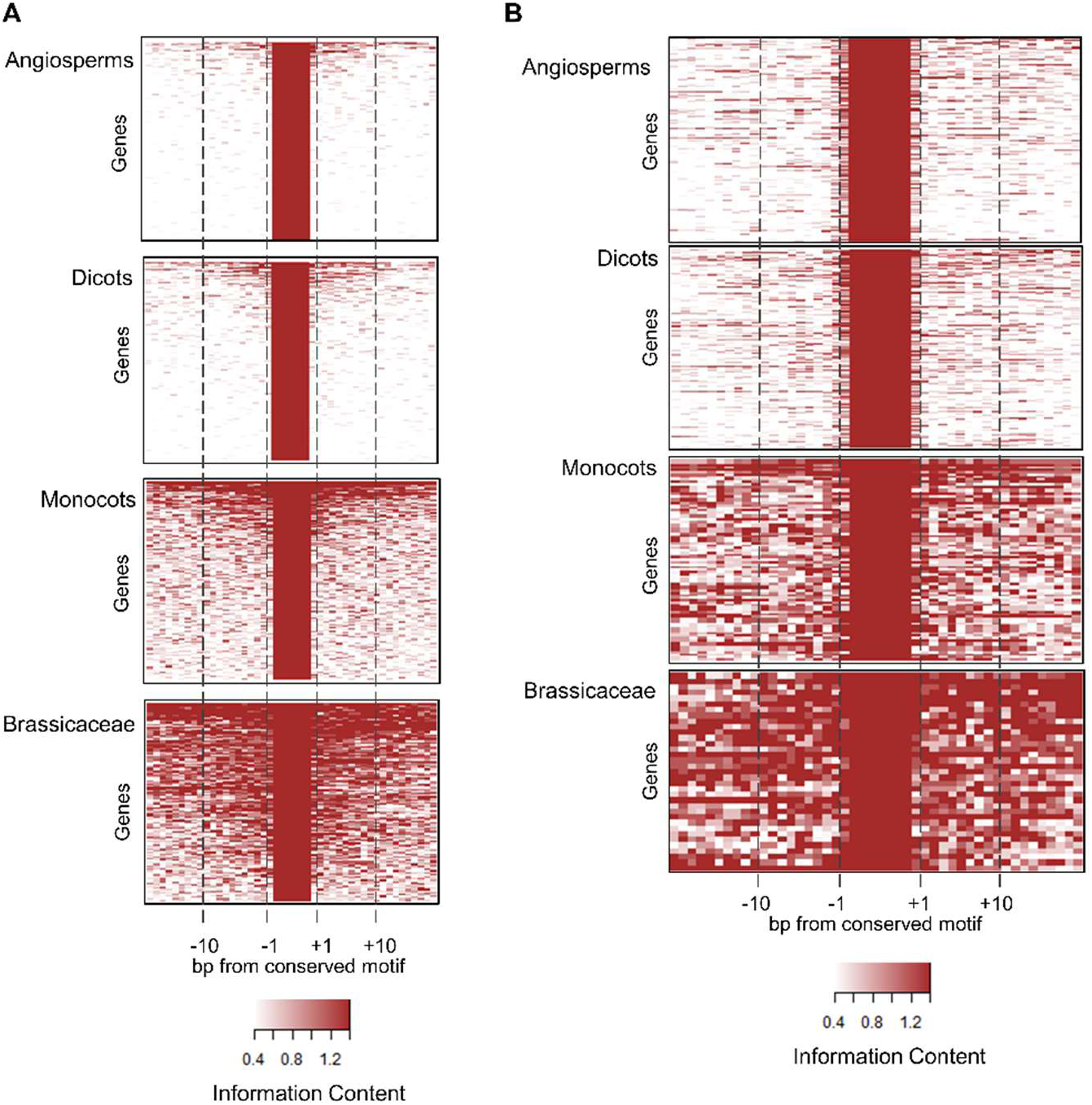
Deep conservation of sequence context for cytRE and abRE variants. **A-B)** Information score values computed from PWM for sequences flanking the conserved cytRE (A) and abRE (B) motifs.

**Supplemental Figure 6.**
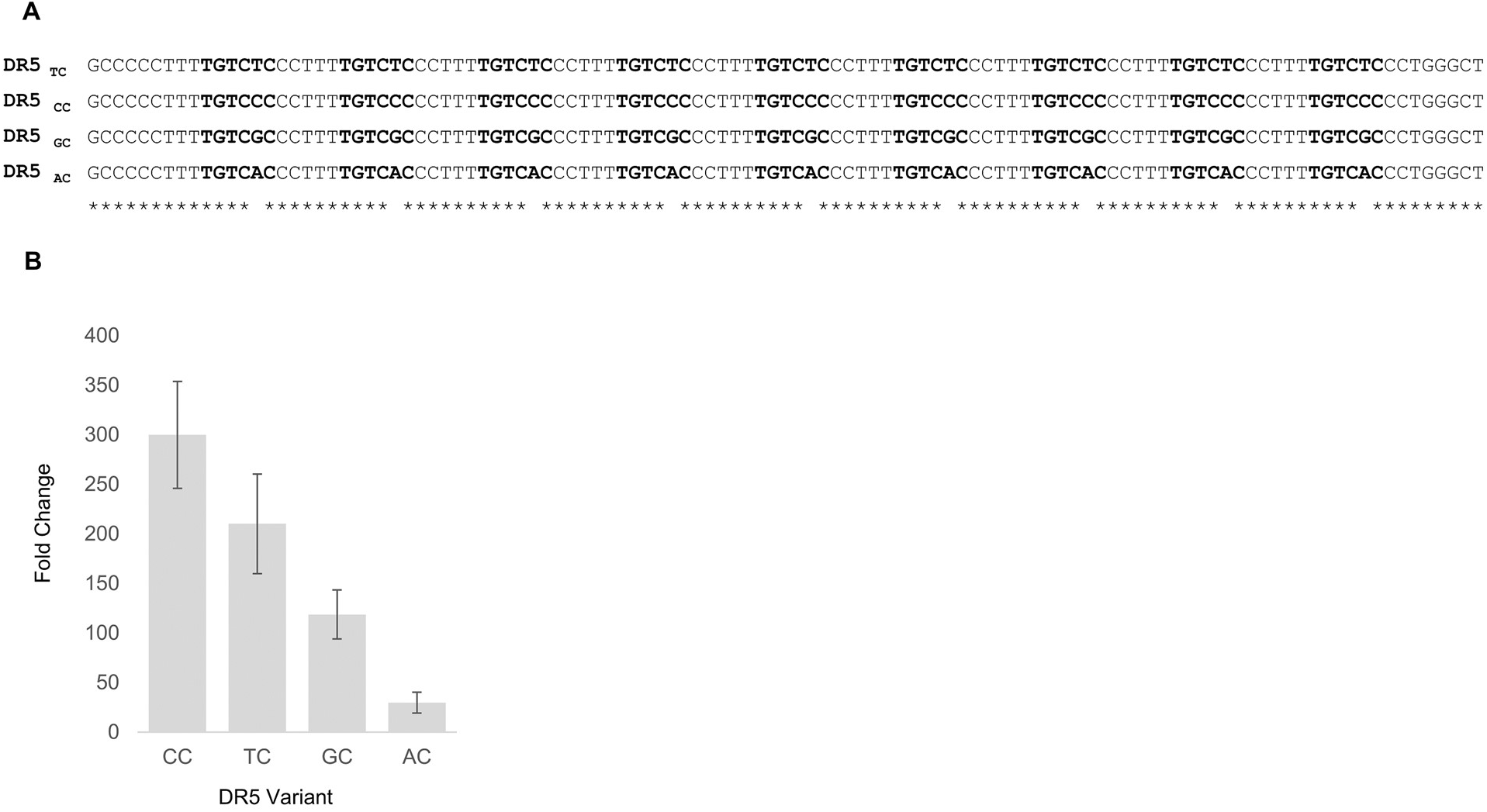
Sequence and response of DR5 variants. **A)** Sequences of the four DR5 variants used in this study. **B)** Fold-change of the GFP transcript 6h following auxin application to roots carrying different DR5 variants, as measure by qPCR. Error bars represent standard error. n=3.

**Supplemental Figure 7.**
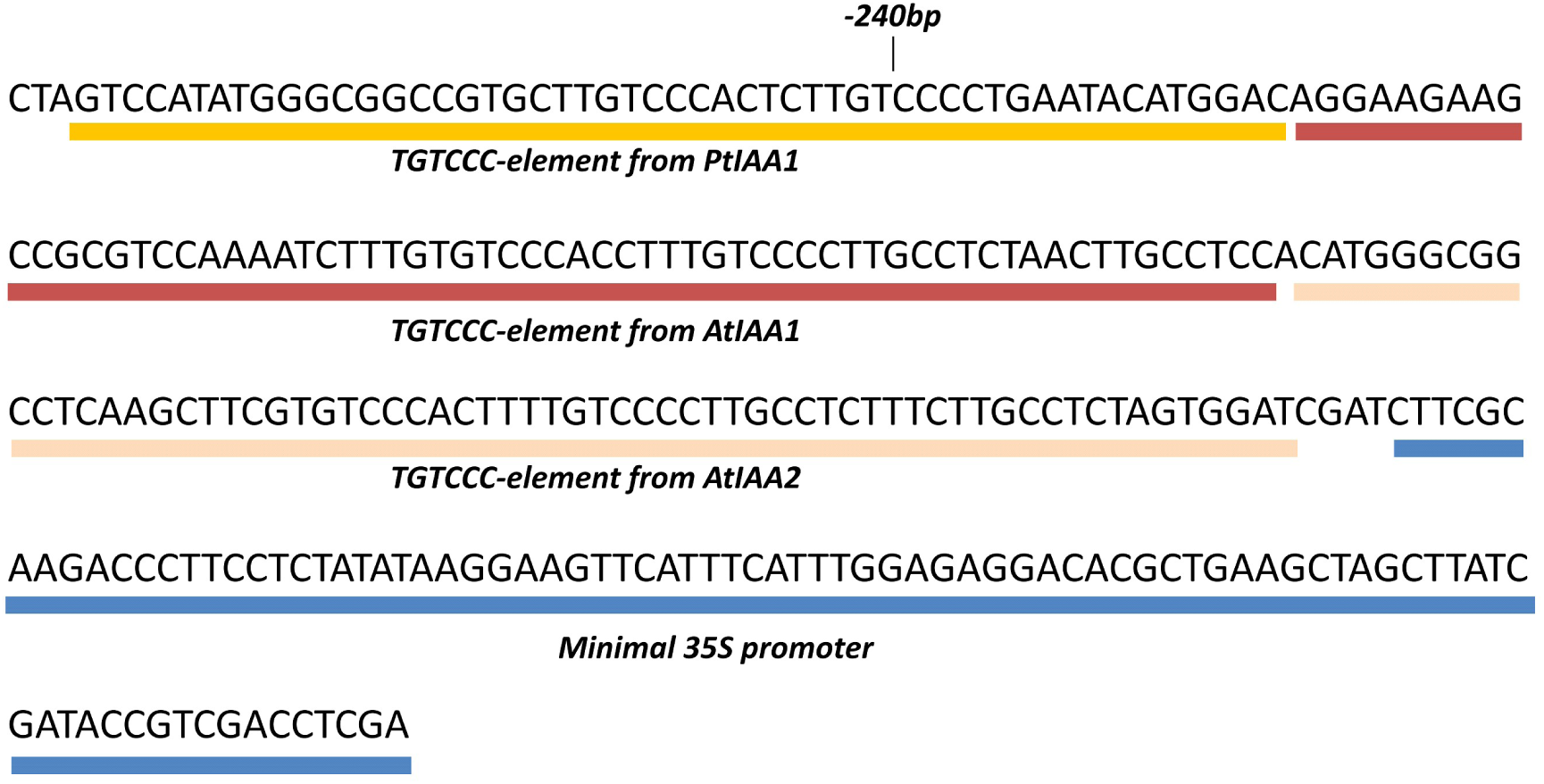
Synthetic pIAA motif high-sensitivity auxin reporter.

**Supplemental Table 1. List of genomes used in this study**

**Supplemental Table 2. Ortholog candidates for all genes in the species used in this study**

**Supplemental Table 3. List of genes with conserved auxRE motifs**

**Supplemental Table 4. List of genes with conserved cytRE motifs**

**Supplemental Table 5. List of genes with conserved abRE motifs**

**Supplemental Table 6. Sequence conservation of sequence flanking conserved auxRE motifs**

**Supplemental Table 7. Sequence conservation of sequence flanking conserved cytRE motifs**

**Supplemental Table 8. Sequence conservation of sequence flanking conserved abRE motifs**

## Material and Methods

### Plant growth and imaging

*Arabidopsis* seeds were planted on ½ Murashige and Skoog agar plates, kept for 48h in the dark at 4°C and transferred to a growth chamber at 21°C in continuous light. Tomatoes were germinated in a growth chamber and moved to a temperature-controlled greenhouse with natural light (26°C/day 18°C/night). Transgenic *Arabidopsis* were generated using floral dipping. Transgenic tomatoes were generated using Agrobacterium (strain GV3103)-mediated cotyledon transformation. Images were taken using a Nikon SMZ18 fluorescence stereoscope, with identical settings for all images. Confocal microscopy was performed using a Leica SP8. For *Arabidopsis* roots propidium Iodide was used to stain cell walls. Tomato lines carrying the pIAAmotif:LhG4 driver were crossed to plants carrying op:mRFP. Tomato roots and adventitious roots were fixed and cleared using ClearSee^51^, cell wall stained was performed using SR2200 (Renaissance Chemicals) prior to mounting and visualization using 405nm, 488nm and 561nm lasers. To image the shoot meristem, live tomato shoots were dissected, mounted in gel and imaged using a Zeiss Lightsheet Z.1.

### Construct generation

For the protoplast assay, DNA for the four DR5 variants was synthesized and cloned, using Gateway, upstream to a minimal 35S promoter fused to Venus. To generate transgenic *Arabidopsis*, the same variants were cloned, using Gateway, to the pKGWFS7 plasmid, upstream to a GFP-GUS fusion protein. For ARF overexpression, the coding sequence was cloned, using Gateway, into a p2GW7,0 plasmid. To generate the pIAA2 and pIAA4 promoter, the sequences 1200bp and 1500bp upstream, respectively, were amplified and cloned into pENTR using Gateway. Site-directed mutagenesis was used to replace the TGTCCC motifs with TGTCAC. The fragments were then shuttled to the pKGWFS7 vector.

The synthetic auxin reporter pIAA motif was constructed by cloning the TGTCCC-containing regulatory regions of the IAA1 ortholog from *Populus trichocarpa*, *IAA1* from *Arabidopsis* and IAA2 from *Arabidopsis*, upstream to a minimal 35S promoter. For *Arabidopsis* lines, the three segments were cloned into a level 0 MoClo part using Golden Gate and then fused to *mNeonGreen-N7* and a heat shock protein terminator to form a level 1 construct. The level 1 was transferred to a level 2, together with a kanamycin resistance cassette. For tomato transformation, the synthetic promoter was cloned upstream to LhG4 using restriction cloning, and then subcloned into pART27.

### Protoplast assay

Arabidopsis plants were grown on agar plates as described above. At 7 days after transfer to light, protoplasts were extracted and transfected with 10µg plasmid carrying the synthetic reporter fused to VENUS, together with a transfection control of 10µg pMon plasmid, carrying 35S:*RFP*, according to (Bargmann et al.^52^). Cell aliquots were treated with 1µM 2,4-D or mock for 6h. Mean YFP and RFP fluorescence of at least 10,000 cells was measured using a BD Accuri 6 plus analyzer. The mean RFP signal was used to normalize the YFP signal. Experiments were performed in triplicates.

### Ortholog calling

For each species, a reciprocal blast was performed between all proteins and the *Arabidopsis* protein sequences using a minimum E-value of 1E-10. Orthologs were ranked based on E-value. An ortholog quality score for each gene was computed as the combined rank in the reciprocal blast, with 2 being the highest score. Gene pairs were sorted based on ortholog quality score and the genes with lowest scores (up to eight) were considered as putative orthologs. The mean number of putative orthologs per species, for each *Arabidopsis* gene was 2.61±0.34 genes.

### Computational identification of motif conservation

Genome sequences and annotation gff3 files were obtained from phytozone 12 and a perl script was used to generate upstream sequences and identify the motifs variants for each gene. Upstream sequences included 500bp upstream and 100bp downstream to the annotated TSS. Calculation of conservation scores were performed as followed. For each *Arabidopsis* gene, the orthologs for all species were obtained, as described above. Of the putative orthologs for a given species, the one with a motif variant composition most similar to the *Arabidopsis* gene was selected as an ortholog. The motif variants for each gene were arrange on a predefined species tree. Maximum parsimony method was used to derive the motif variants on the internal nodes. The conservation score for each motif variant was calculated as the number of occurrences of the variant in the tree minus the number of changes (gain or loss) this variant had across the tree. To determine the background distribution, conservation scores of nucleotides at positions 8bp away from the core motif, were calculated for all genes. A negative binomial distribution was fitted to the overall conservation score for all variants at position - 8bp. This fitted distribution was used to compute p-values for the conservation of the motif variants. Significance cutoffs for the –log2(pval) were set to 23, 21 and 35 for the auxRE, abRE and cytRE motifs, respectively. The R source code is deposited at https://github.com/idanefroni/CoMoVa.

### GO term analysis

GO term analysis was performed for the *Arabidopsis* orthologs of the genes with conserved motifs, using the topGO package with the “weight01” algorithm. P-values were calculated used the hypergeometric test without further correction to multiple testing. Enrichment testing was performed for each variant individually and then for the entire list of genes with conserved variants.

### Identification of auxin responsive genes

For the root response, tomato seedlings were germinated on ½ Murashige and Skoog plates and grown in a growth chamber at 22C under continuous light. 4 days after germination, were transferred to plates containing 1µM of the auxin analog 2,4-D (Sigma D7299) for 3h, followed by excision of 1cm root tips and flash freezing in liquid nitrogen. For shoot samples, three-week-old M82 plants were sprayed with either mock or 1 µM of the auxin analog Picloram (Sigma P5575) for 3 hours and 15 young leaf primordia (Leaf number 5 at the P5 stage) were collected and flash frozen. RNA-extraction was performed using Qiagen RNAEasy Microkit and sequencing libraries prepared using Lexigen 3’ Quant-Seq kit, according to the manufacturer instructions. Experiments were performed in duplicates or triplicates. Single-end sequencing was performed using Illumina Nextseq 500. Expression calling was performed using Salmon 0.8.2, by aligning to the tomato genome ITAG3.2. As the library is 3’ biased, we extended all genes to cover 500 additional base pairs downstream. Expression data were normalized using Deseq2 and genes induced at False Discovery Rate (FDR)<0.01 were selected. Tomato auxin response data was submitted to GEO (GSExxx). Processed expression values for the *Arabidopsis* and *Brachypodium* root auxin response were obtained from previously published analysis^7,35^. Maize auxin response data was obtained from GEO (GSE111792), aligned to the maize transcriptome (AGPv3, obtained from Phytozone) and auxin responsive genes were identified as above.

